# Development of a free radical scavenging probiotic to mitigate coral bleaching

**DOI:** 10.1101/2020.07.02.185645

**Authors:** Ashley M. Dungan, Dieter Bulach, Heyu Lin, Madeleine J. H. van Oppen, Linda L. Blackall

## Abstract

Corals are colonized by symbiotic microorganisms that exert a profound influence on the animal’s health. One noted symbiont is a single-celled alga (from the family *Symbiodiniaceae*), which provides the coral with most of its fixed carbon. During thermal stress, hyperactivity of photosynthesis results in a toxic accumulation of reactive oxygen species (ROS). If not scavenged by the antioxidant network, ROS may trigger a signaling cascade ending with the coral host and algal symbiont disassociating; this process is known as bleaching. Our goal was to construct a probiotic comprised of host-associated bacteria able to neutralize free radicals such as ROS. Using the coral model, the anemone *Exaiptasia diaphana*, and pure bacterial cultures isolated from the model animal, we identified six strains with high free radical scavenging ability belonging to the families *Alteromonadaceae, Rhodobacteraceae, Flavobacteriaceae*, and *Micrococcaceae*. In parallel, we established a “negative” probiotic consisting of genetically related strains with poor free radical scavenging capacities. From their whole genome sequences, we explored genes of interest that may contribute to their potential beneficial roles, which may help facilitate the therapeutic application of a bacterial probiotic. In particular, the occurrence of key pathways that are known to influence ROS in each of the strains has been inferred from the genomes sequences and are reported here.

**IMPORTANCE:** Coral bleaching is tightly linked to the production of reactive oxygen species (ROS), which accumulates to a toxic level in algae-harboring host cells leading to coral-algal dissociation. Interventions targeting ROS accumulation, such as the application of exogenous antioxidants, have shown promise for maintaining the coral-algal partnership. With the feasibility of administering antioxidants directly to corals being low, we aim to develop a probiotic to neutralize toxic ROS during a thermal stress event. This probiotic can be tested with corals or a coral model to assess its efficacy in improving coral resistance to environmental stress.

## INTRODUCTION

Coral reefs are among the most biologically and economically valuable ecosystems on Earth (1-3). While they cover less than 0.1% of the ocean floor (4), coral reefs support fisheries, tourism, pharmaceuticals and coastal development with a global value of $8.9 trillion “international $”/year (5). Corals and other reef organisms have been dying, largely due to anthropogenic influences such as climate change (6, 7), which has led to an increased frequency, intensity and duration of summer heat waves that cause coral bleaching (8, 9).

The coral holobiont, which is the sum of the coral animal and its associated microbiota, including algae, fungi, protozoans, bacteria, archaea and viruses (10), is an ecosystem engineer. By secreting a calcium carbonate skeleton, the reef structure rises from the ocean floor, forming the literal foundation of the coral reef ecosystem. The success of corals to survive and build up reefs over thousands of years (11) is tightly linked to their obligate yet fragile symbiosis with endosymbiotic dinoflagellates of the family *Symbiodiniaceae* (12).

Intracellular *Symbiodiniaceae* translocate photosynthetically fixed carbon to the coral host (13, 14) in exchange for inorganic nutrients and location in a high light environment with protection from herbivory (15, 16). During periods of thermal stress, the relationship between the coral host and their *Symbiodiniaceae* can break down, resulting in a separation of the partners and significantly, fixed carbon shortage for the host. This phenomenon, ‘coral bleaching’, is devastating to the host and detrimental to the reef system. The ecosystem-wide effects of bleaching on the coral include reduced skeletal growth and reproductive activity, a lowered capacity to shed sediments, and an inability to resist invasion of competing species and diseases. Severe and prolonged bleaching can cause partial to total colony death, resulting in diminished reef growth, the transformation of reef-building communities to alternate, non-reef building community types, bioerosion and ultimately the disappearance of reef structures (12).

There are several hypotheses detailing the mechanisms driving bleaching (see 17, 18-20), with a common theme being the overproduction and toxic accumulation of reactive oxygen species (ROS) from the algal symbiont. Excess ROS are generated by a number of pathways including heat damage to both chloroplast and mitochondrial membranes (21, 22), and are shown to play a central role in inter-partner communication of a stress response (17). Once generated, ROS causes damage to many cell components including photosystem II reaction centers in the *Symbiodiniaceae*, specifically at the D1 and D2 proteins (23, 24), thus interfering with the supply of fixed carbon to the holobiont. Once damaged, *Symbiodiniaceae* are no longer able to maintain their role in the symbiotic relationship with corals and separate from the host tissue via *in situ* degradation, exocytosis, host cell detachment, host cell apoptosis or host cell necrosis (17).

Probiotics are preparations of viable microorganisms that are introduced to a host to alter their microbial community in a way that is beneficial to the system. Microbiome engineering through the addition of probiotics has been postulated as a key strategy to facilitate adaptation to changing environmental conditions by enhancing corals with the metabolic capabilities of the introduced probiotic bacterial strains (25-30). The differences in the bacterial species composition of healthy and thermally stressed corals (31-36) and the coral model *Exaiptasia diaphana* (37-39) suggest a role for microbiome engineering in cnidarian health. A disruption to the bacterial community of *Pocillopora damicornis* with antibiotic treatment diminished the resilience of the holobiont during thermal stress, whereas intact microbial communities conferred resilience to thermal stress and increased the rate of holobiont recovery after bleaching events (40). The relative stability of coral-associated bacterial communities has also been linked to coral heat tolerance; for instance, the bacterial community of heat sensitive *Acropora hyacinthus* corals shifted when transplanted to thermal stress conditions, whereas heat-tolerant *A. hyacinthus* corals harbored a stable bacterial community (41).

In recent years, researchers have begun to explore the use of probiotics in corals and the model organism for corals, *E. diaphana*. To inhibit the progression of white pox disease, caused by pathogenic *Serratia marcescens*, an *Alphaproteobacteria* cocktail containing several *Marinobacter* spp. isolates was applied to *E. diaphana* (42). These introduced strains were able to inhibit both biofilm formation and swarming of *S. marcescens*, which halted disease progression. The *Marinobacter*-based probiotic was deemed effective as anemones exposed to both the cocktail and pathogen survived after seven days, while anemones in the *S. marcescens* control treatment died. A bacterial consortium native to the coral *Mussismilia harttii* was selected to degrade water-soluble oil fractions(43). This bioremediation strategy reduced the negative impacts of oil on *M. harttii* health and accelerated the degradation of petroleum hydrocarbons (43). Coral microbiomes have also been manipulated through addition of a consortium of native or seawater derived bacteria to the surface of *P. damicornis* to mitigate the effects of thermal stress (44). The results from this study suggest the consortium was able to partially mitigate coral bleaching.

Our goal was to identify bacterial strains suitable for use in a probiotic to mitigate the effects of thermal stress in *E. diaphana*. Given the potential role of ROS in the bleaching process, our focus was to select diverse *E. diaphana*-sourced bacterial strains with antioxidant properties while avoiding potential pathogens. Antioxidant properties were measured using the stable free radical 2,2-diphenyl-1-picrylhydrazyl (DPPH), which is reduced in the presence of an antioxidant molecule, undergoing a color change from a violet to a colorless solution.

## RESULTS

### Diversity of culturable bacteria associated with *E. diaphana*

A total of 842 isolates were obtained from four genotypes of Great Barrier Reef (GBR)-sourced *E. diaphana*. There were no significant differences in bacterial colony forming units (CFUs) between the four genotypes, regardless of growth medium, with 5.9-10.3 × 10^3^ cells per anemone on Reasoner’s 2A agar (R2A) and 6.3-10.4 x10^3^ cells per anemone on marine agar (MA) (p>0.05). Partial 16S rRNA gene sequences (∼1000 bp) were used to identify the closest matches from the NCBI database using the Basic Local Alignment Search Tool (BLASTn). In total there were 109 species in 64 genera, 27 families and six phyla (Fig. 1). The most abundant genera were *Alteromonas, Labrenzia*, and *Ruegeria* (Table 1). Gram-positive bacteria comprised 23 species, including *Microbacterium* (31 isolates) and *Micrococcus* (28 isolates). Eight genera were found to be associated with all four genotypes (Table 1); these eight genera made up 59.4% of all *E. diaphana*-associated bacterial isolates.

**Table 1:** Bacterial genera associated with all four genotypes of GBR-sourced *E. diaphana* (AIMS1-4).

| Genus | Class | No. of species | AIMS1 | AIMS2 | AIMS3 | AIMS4 | *Total Isolates |
| --- | --- | --- | --- | --- | --- | --- | --- |
| ***Alteromonas*** | *Gamma-proteobacteria* | 6 | 10 | 52 | 24 | 30 | 116 |
| ***Labrenzia*** | *Alpha-proteobacteria* | 4 | 11 | 10 | 26 | 38 | 85 |
| ***Marinobacter*** | *Gamma-proteobacteria* | 4 | 7 | 25 | 10 | 13 | 55 |
| ***Muricauda*** | *Flavobacteriia* | 1 | 16 | 11 | 10 | 5 | 42 |
| ***Roseovarius*** | *Alpha-proteobacteria* | 3 | 9 | 12 | 5 | 6 | 32 |
| ***Ruegeria*** | *Alpha-proteobacteria* | 3 | 29 | 8 | 5 | 39 | 81 |
| ***Shimia*** | *Alpha-proteobacteria* | 2 | 2 | 3 | 9 | 40 | 54 |
| ***Vibrio*** | *Gamma-proteobacteria* | 3 | 3 | 7 | 31 | 15 | 56 |

\*521 out of the 842 isolates obtained from the four *E. diaphana* genotypes sourced from the Great Barrier Reef.

**FIG 1:**
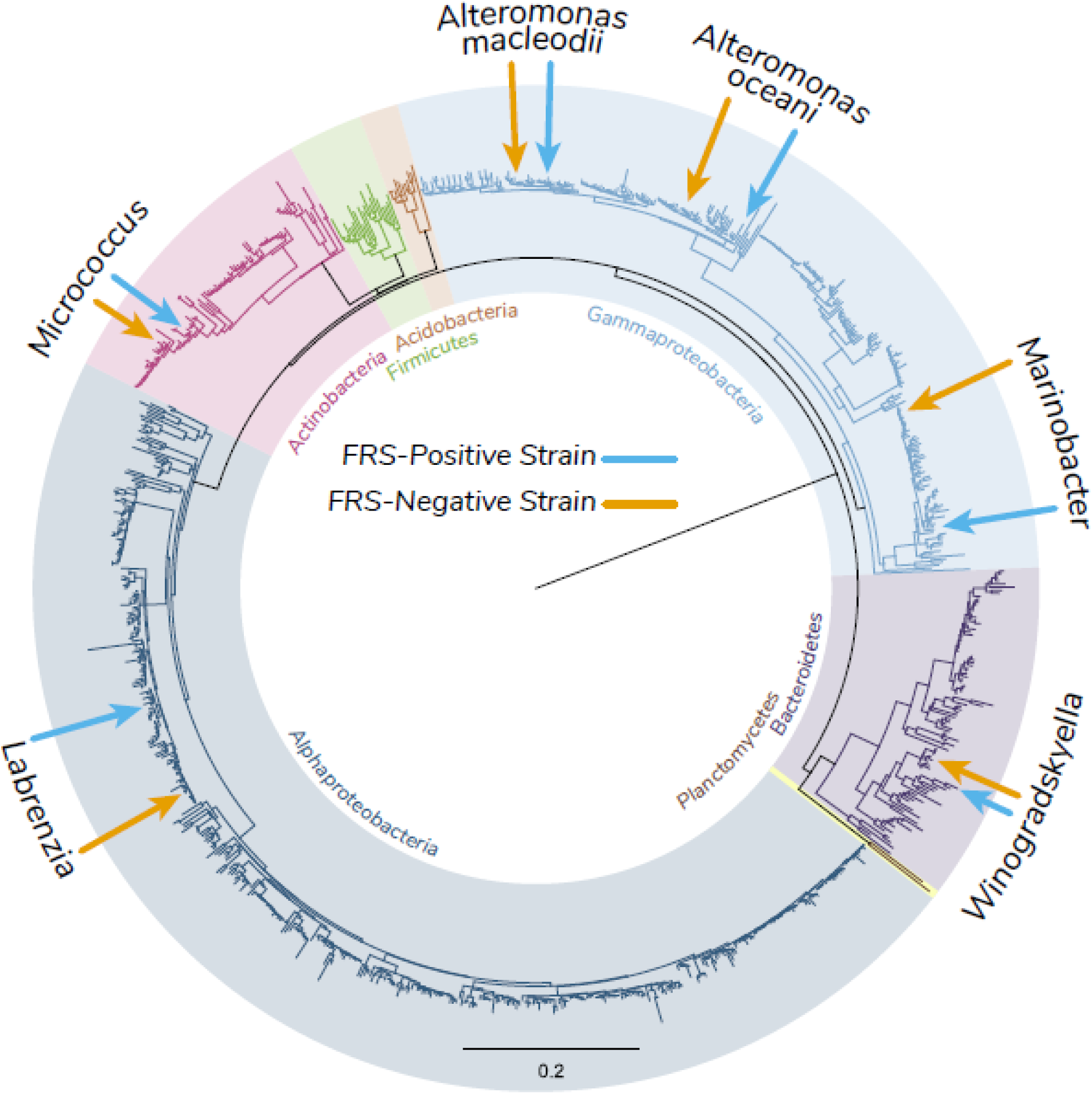
Neighbor-Joining tree showing an overview of the phylogenetic relationship between the 842 *E. diaphana*-associated bacterial isolates inferred using partial 16S rRNA sequences. These isolates covered six phyla indicated by shading over the tree with *Proteobacteria* split into the classes *Gammaproteobacteria* and *Alphaproteobacteria*. The positions of selected probiotic strains are highlighted by arrows with blue arrows indicating the high FRS strains and orange arrows indicating low FRS strains.

### Bacterial probiotic selection

Of the original 842 isolates, 709 were screened for their ability to scavenge free radicals. Those isolates were divided into three categories, positive (144), weakly positive (121), and negative (444). Ninety-eight strains representing eight families and 18 genera were then quantitatively assessed for FRS. There was no clear pattern of free radical scavenging (FRS) capacity at the family level (Fig. 2) with strain-specific responses evident. From these isolates, probiotic members were selected by choosing *E. diaphana*-associated bacteria, where conspecific or congeneric pairs of strains displayed a high (“positive”) and low (“negative”) FRS ability (Fig. 2; Table 2). Of the 12 selected probiotic members, seven were catalase positive and five were catalase negative (Table 2). In each probiotic set (i.e., high or low FRS strains), none of the selected isolates showed antagonistic activity against one other as evidenced by the absence of any zone of inhibition and growth from each combination of isolates on a plate.

**Table 2:**
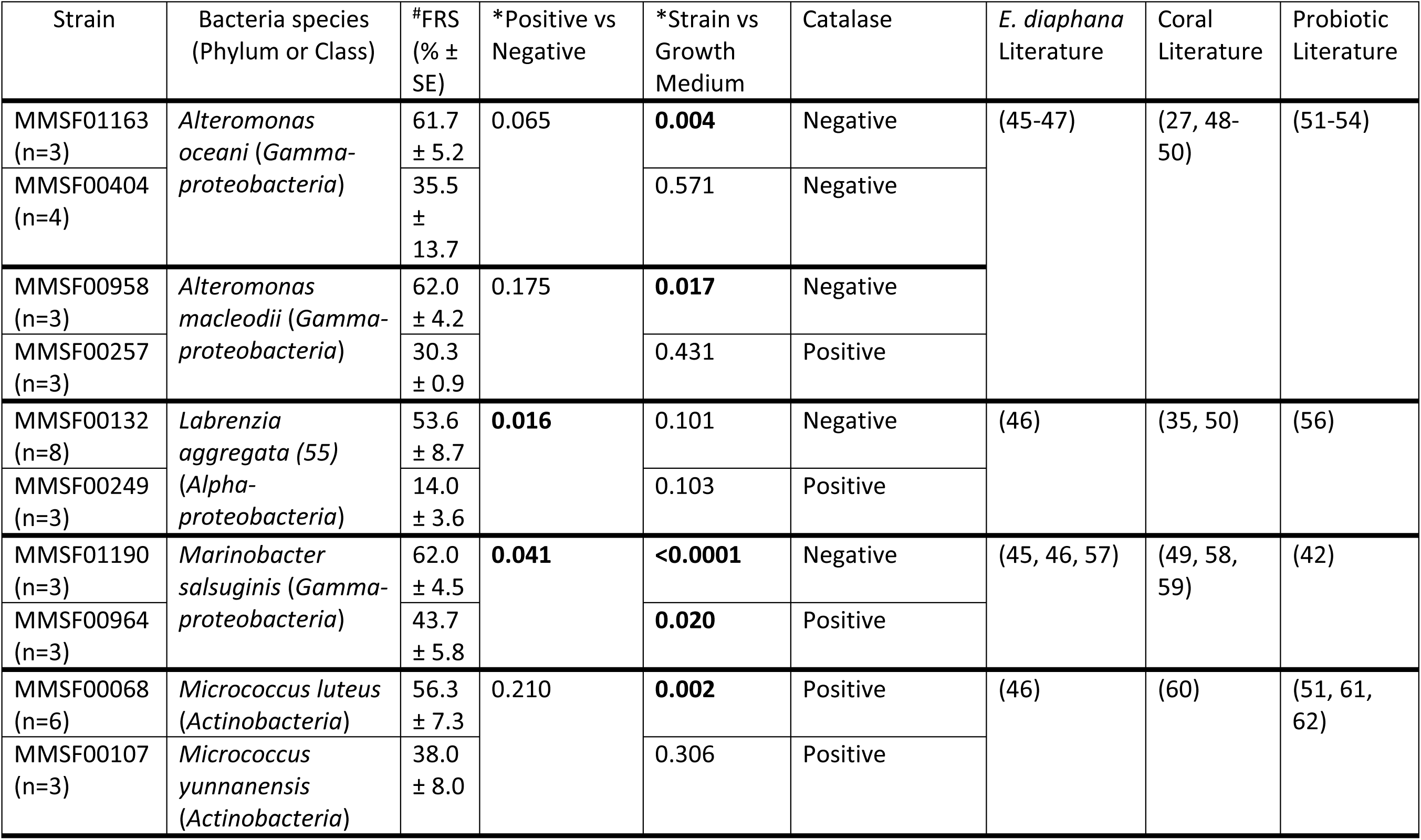
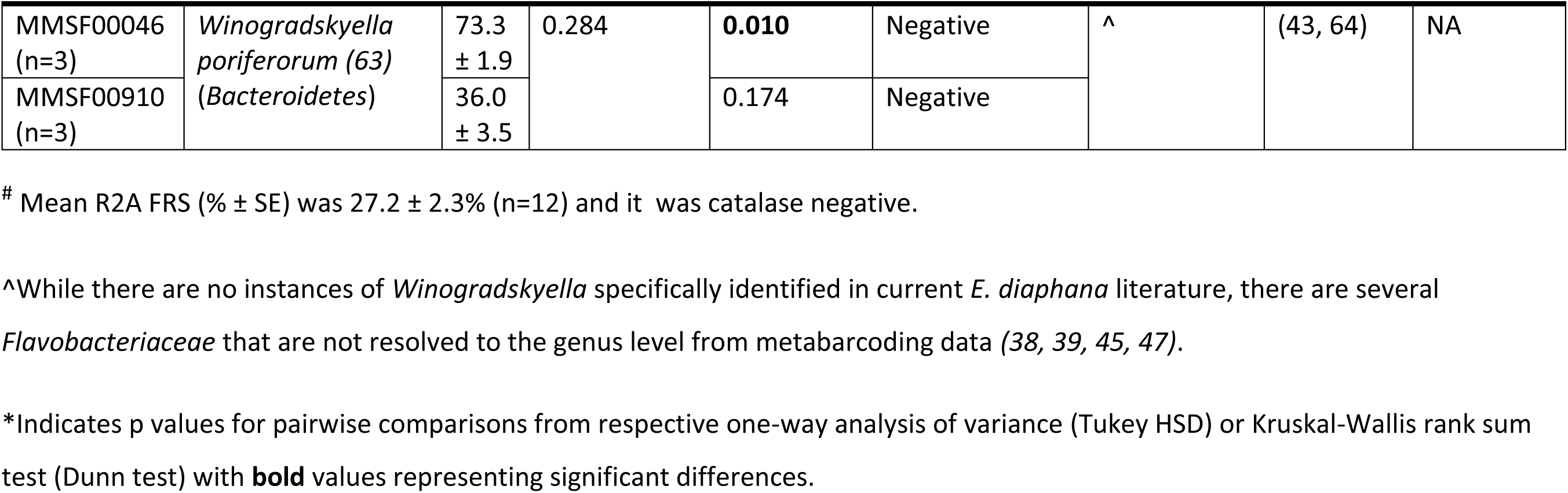
Overview of probiotic high FRS bacterial strains and conspecific/congeneric low FRS control strains. All sequence data can be found under BioProject# PRJNA574193. References to each probiotic candidate at the Genus level are identified in the last three columns.

**FIG 2:**
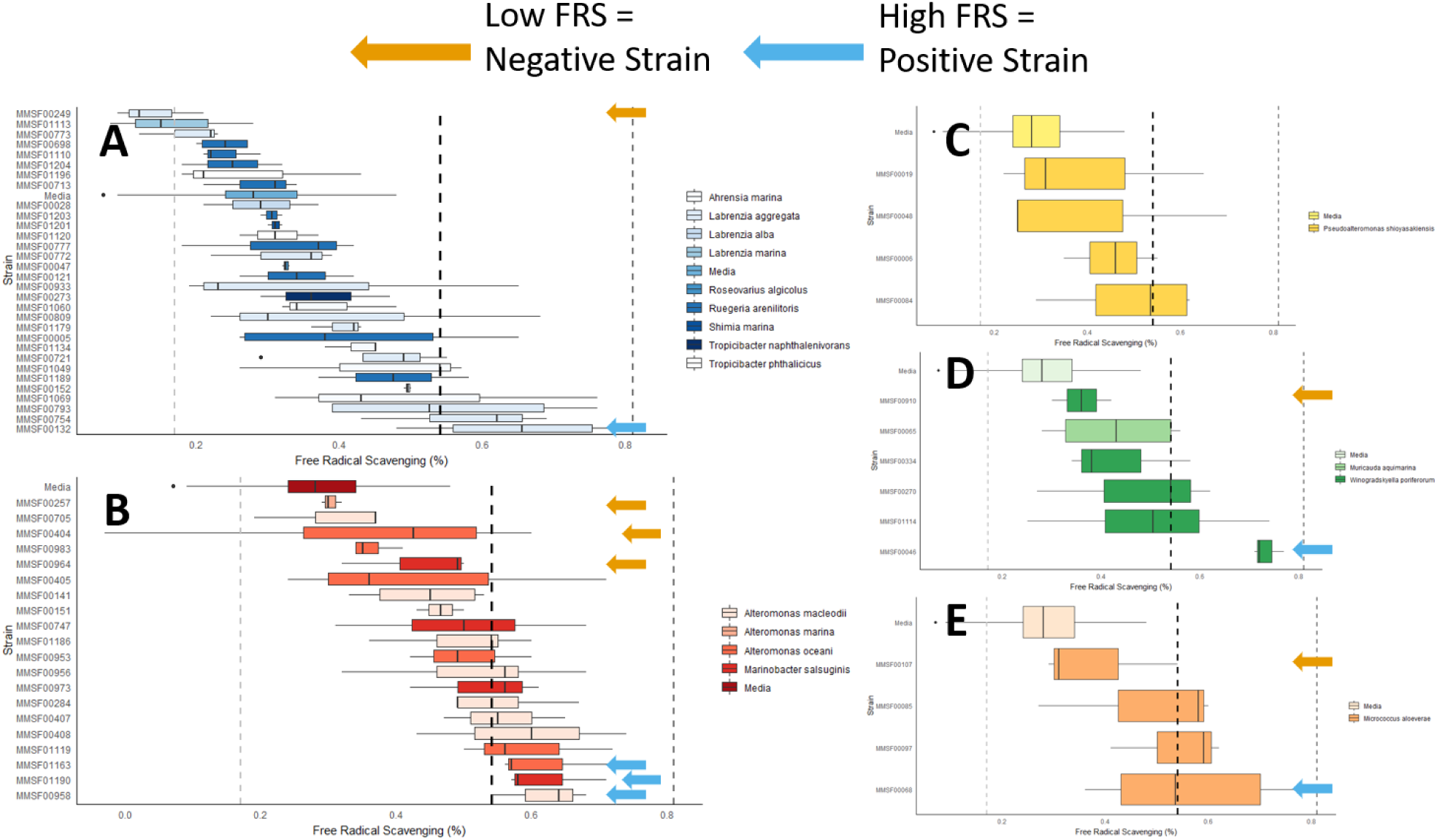
Quantitative FRS ability of *E. diaphana*-associated bacteria isolates, separated by Family. Families with high relative abundance among all cultured bacteria (*Rhodobacteraceae* – A, *Alteromondaceae* – B, *Pseudoalteromonadaceae* – C, *Flavobacteraceae* – D, and *Micrococcaceae* – E) were separately analyzed to identify strains with a high FRS ability (blue arrows) and a corresponding conspecific or congeneric strain with a low FRS ability (orange arrows). In each panel, the light dashed vertical line on the left represents the mean FRS of a 0.025% (w/v) ascorbic acid standard, the middle dark dashed vertical line is the mean FRS for 0.05% (w/v) ascorbic acid standard, and the far right dashed line is the mean FRS of the 0.075% (w/v) ascorbic acid standard.

### Comparative genomics

As part of the characterization of the 12 isolates (six positive and six conspecific or congeneric negative FRS isolates), a draft genome sequence was determined for each isolate. A summary of the data and metrics for the draft genome sequences is presented in Table S1. The diversity of the six pairs of isolates is indicated by the %G+C range (35% to 72%) and genome size (2.4 Mb to 6.8 Mb). Each isolate pair was classified as the same species except the *Micrococcus* stains. Isolates MMSF00068 (high FRS strain) and MMSF00107 (low FRS strain) are classified as *Micrococcus luteus* by both 16S rRNA gene and genome-based methods, but MMSF00107 is classified as *Micrococcus yunnanensis* by NCBI. These isolates have the smallest genomes among the isolates (2.43 Mb and 2.48 Mb, respectively) and the highest G+C content of 72.8% and 72.4%, respectively. The classification of isolate MMSF00107 is uncertain, particularly given that the genome sequence for the type strain for *M. yunnanensis* is not yet available.

Core genome comparison of the draft genome sequences provides an overview of the genetic relationship between the conspecific or congeneric isolate pairs. A wide range of genome variation between pairs was observed, this ranged from ∼190,000 core genome single nucleotide polymorphisms (SNP) differences between the *Alteromonas oceani* strains, to fewer than five core genome SNPs between the *Labrenzia aggregata* isolates (Table S2). Importantly, this is not a comprehensive estimate of isolate identity, e.g. the genome sequences of *Winogradskyella poriferorum* isolates MMSF00910 and MMSF00046 are nearly identical with fewer than ten pairwise SNP differences in the core genome; but there are accessory genome differences.

### Genes of interest

The annotated genome sequences of each selected probiotic member were searched for key genes of interest (Table S3-S4). Dimethylsulfoniopropionate (DMSP) cleavage to dimethylsulfide (DMS) was identified by presence of one or more of the DMSP lyase genes; *dddP, dddD, dddL, dddW*, and *dddQ*. Only *L. aggregata* strains contained DMSP lyase genes (*dddP* and *dddL*) in their whole genome sequences. DMSP biosynthesis was identified by the presence of *dsyB*, which is the only described gene for an enzyme in the DMSP biosynthesis pathway. The *cobP* gene was used as an indicator for the presence of the dynamic vitamin B_12_ pathway, which contains 27 genes (Table S4). Again, only the *L. aggregata* isolates contained *cobP*. Catalase positive strains were identified by the presence of *katG*; all positive and negative FRS strains contained *katG* except the *Micrococcus* spp. strains, in which *katA* and *katE* were detected.

### 16S rRNA gene copy number

The 16S rRNA gene copy numbers of the 12 draft genomes were estimated using a read depth approach (Table S1). The copy numbers were similar within pairs of isolates, in which the pair of isolates MMSF00257 and MMSF00958 contained the most copies (5.15 and 4.79, respectively), corresponding with copy numbers in a known closed genome of *A. macleodii* (65). *Marinobacter* isolates MMSF00046 and MMSF00910 contained the fewest copies (1.03 and 0.77, respectively).

## DISCUSSION

The 842 *E. diaphana* bacterial isolates reported here comprise 109 species from 64 genera and six phyla. Using metabarcoding, studies of microbiomes associated with *E. diaphana* have revealed a similar diversity at the phylum level for *E. diaphana* sourced from the GBR (37), Hawaii (strain H2; 45), Pacific and Caribbean (57), Atlantic (strain CC7; 47) and Red Sea (38), as well as stony corals (66). Thus, our culture collection of *E. diaphana* bacterial isolates captures the diversity of the *E. diaphana*-associated microbiome. The *E. diaphana* probiotics generated in this study were assembled from 12 isolated bacterial strains selected based on their FRS ability since free radical production, specifically ROS, is relevant in coral bleaching. The broader culture collection contains bacteria with a wide range of FRS capacity, and our probiotic comprises greater bacterial diversity compared to others (42, 44).

The consistent and frequent reporting of our probiotic bacteria genera in *E. diaphana* and coral studies (Table 2) suggests these bacteria likely have key functions in cnidarian holobionts. Among these potential functions are the production of the antioxidant DMSP and the breakdown of DMSP to other antioxidants (67). *L. aggregata* has been reported to produce DMSP in the absence of any methylated sulfur compounds with *dsyB* identified as the first DMSP biosynthesis gene in any organism (68). The *dsyB* gene was found in the genome sequences of both high and low FRS *L. aggregata* strains (Table S3). Many *E. diaphana*-sourced bacterial species, specifically relatives of our selected probiotic members, are implicated in the degradation of DMSP to DMS (*Alteromonas* spp., (69); *Labrenzia* spp., (70)). *dddP* codes for the enzyme responsible for cleaving DMSP to DMS and acrylate and was used (from the Prokka annotation) as an indicator of a DMSP degradation genotype. Only the *L. aggregata* isolates contained genes responsible for DMSP degradation (Table S3).

Carotenoids are among the strongest antioxidants and are highly reactive against both ROS and free radicals (71-75). Carotenoids are lipid-soluble pigments, and in bacteria they give an orange-yellow hue to colonies. Two of the five selected probiotic genera produce orange/yellow colonies (*Winogradskyella, Micrococcus*), and there is evidence of carotenoid production by marine *Flavobacteriaceae* (75, 76) and *Micrococcus* (77). A marine *Flavobacteriaceae* (strain GF1) was found to produce the potent antioxidant carotenoid zeaxanthin that protected *Symbiodiniaceae* from thermal and light stress (78).

Vitamin B_12_ is a cofactor involved in the production of the amino acid methionine, which is needed to synthesize every protein and in diverse metabolic pathways including generation of the antioxidants glutathione and DMSP (79). Vitamin B_12_ is synthesized by many heterotrophic bacteria (80). Genomic evidence suggests that *Symbiodiniaceae* have lost the capacity to synthesize vitamin B_12_ (81), which is in agreement with other work showing that free-living *Symbiodiniaceae* rely on bacterial symbionts for this important cofactor (82). The genes involved in the biosynthesis of vitamin B_12_ have been found in coral-associated bacteria, specifically *L. aggregata* cultured from the Caribbean coral, *Orbicella faveolata* (83). Eighteen genes, including the *cobP* gene (80), are annotated in each of *L. aggregata* isolates (MMSF00132 & MMSF00249), suggesting both are capable of vitamin B_12_ biosynthesis. None of the other sequenced isolated had genes related to vitamin B_12_ biosynthesis, except in the *Marinobacter salsuginis* isolates, where a *cobO* gene was detected, strongly indicating that none of the remaining ten isolates were capable of vitamin B_12_ biosynthesis (Table S4).

Bacteria have developed highly specific mechanisms to protect themselves against oxidative stress, in particular using enzymatic antioxidants such as catalase/peroxidase and superoxide dismutase (84). It has been suggested that increasing the *in hospite* concentration of catalase in the coral holobiont by the application of a probiotic with catalase-positive organisms, could possibly minimize the impact of thermal stress by neutralizing hydrogen peroxide (H_2_O_2_) (29). Here we tested all probiotic candidates for catalase production using a standard H_2_O_2_ assay (85). Catalase participates in cellular antioxidant defense by enzymatically breaking down H_2_O_2_ to H_2_O and O_2_. Hydroperoxidase I (HPI, *katG*) is present during aerobic growth and is transcriptionally controlled at different levels, and hydroperoxidase II (HPII, *katE*) is induced during stationary phase (86, 87). The phenotypically determined catalase positive strains (MMSF00249, *L. aggregata*; MMSF00257, *A. macleodii*; MMSF00964, *M. salsuginis*) had a catalase-peroxidase gene (HPI, *katG*); but all strains had at least one *katG, katA* (catalase), or *katE* (catalase HPII) gene (Table S3).The lack of correlation between genotype and phenotype in relation to catalase activity may be associated with the culture conditions used or the bacterial growth phase during the catalase assay. The inconsistency between the catalase and DPPH phenotype and FRS ability (in many cases high FRS isolates were catalase negative while the low FRS strain was catalase positive), justifies our approach of using FRS analysis, instead of using the presence of catalase genes as a proxy for FRS.

A critical characteristic in the selection of probiotic members is the maintenance and proliferation of the inoculated bacteria in the host over time (51) and their potential transmission to the next generation. There is evidence that corals release bacteria with their offspring such as *Alteromonas* (48), *Flavobacteriaceae* (48), *Rhodobacteraceae* (88), and *Marinobacter* (58). While broadcast spawning corals do not appear to transfer their bacterial symbionts with their gametes (vertical transfer) (89), the brooding coral *Porites astreoides* transmits some bacteria vertically to planulae with two bacterial taxa (*Roseobacter* clade-associated bacteria and *Marinobacter* spp.) consistently and stably associating with juvenile *P. astreoides* (58). *Marinobacter* potentially has antioxidant properties (90), while *Roseobacter* spp. might be beneficial in facilitating larval settlement. If adult corals stably associate with inoculated probiotic candidates like *Marinobacter, Alteromonas*, and *Winogradskyella*, they may be passed on to offspring and thus have a long-term positive impact on these individuals.

The probiotic members were chosen from a highly diverse pool of *E. diaphana*-sourced bacterial isolates. While the selected probiotic members are phylogenetically diverse, potentially promising probiotic bacteria in the culture collection were omitted based on our selection criteria. For example, *Ruegeria* spp. were reported to breakdown DMSP and participate in denitrification (83). *Muricauda* isolates had high FRS abilities, but they did not grow consistently in the selected medium and therefore were excluded from the consortium. *Muricauda* isolates have genes for denitrification (83), can breakdown DMSP (70), and produce potent carotenoids (91) that can mitigate thermal and light stresses in *Symbiodiniaceae* cultures (78). *Muricauda* will be included in future probiotic evaluations.

Interactions among members of the microbiota associated with marine animals are undoubtedly complex. Results presented in this manuscript show that pure cultured bacteria from *E. diaphana* can scavenge free radicals, albeit at a strain-specific rate. Inoculation with high FRS strains could be beneficial to the host under high oxidative stress conditions, such as those that contribute to coral bleaching. Outlined here is the start of a complex process to identify, evaluate, and select durable and useful probiotics that can buffer the coral host against climactic variation. Conspecific or congeneric pairs of bacteria provide an opportunity to determine the genetic basis for measured phenotypic differences between the pairs. An essential element of future work will be to investigate the stability of the phenotypic differences observed and this stability may be reflected in the nature of the genetic differences between the pairs of strains.

## MATERIALS AND METHODS

### Isolation of bacterial isolates

GBR origin *E. diaphana* were maintained in the laboratory at 26 °C (92) and used to isolate probiotic candidates. Sixteen individuals from each of four *E. diaphana* genotypes (AIMS1-4) were collected using sterile disposable pipets and gently transferred to filter-sterilized (0.2 µm) reverse osmosis (RO) water reconstituted Red Sea Salt(tm) (Red Sea; RSS) at ∼34 parts per thousand (ppt) salinity (fRSS). This was done to remove some of the influence of the external seawater on the bacterial community. After 30 min, each anemone was transferred to a sterile glass homogenizer with 1 mL of fRSS. Each homogenate was used to prepare serial dilutions from 10^−1^ to 10^−4^. From each dilution, 50 µL was spread plated onto three replicate plates each of MA (Difco(tm) Marine Agar 2216) and R2A (CM0906, Oxoid) supplemented with 40 g L^−1^ RSS and incubated at 26 °C. After one week, CFU counts were completed. Individual bacterial isolates were sub-cultured to purification from plates with <100 CFUs onto the initial isolation medium. All purified bacterial isolates were resuspended in 40% glycerol, aliquoted into 1.2 mL cryotubes and stored at −80 °C.

### Identification of *E. diaphana*-sourced isolates

Colony PCR with the universal bacterial primers 27f (5’ – AGA GTT TGA TCM TGG CTC AG – 3’) and 1492r (5’ – TAC GGY TAC CTT GTT ACG ACT T – 3’) (93) was used to generate 16S rRNA gene amplicons from each isolate. Briefly, cells from each pure culture were suspended in 20 µL Milli-Q water and denatured at 95 °C for 10 min. The suspension was then centrifuged at 2,000 x *g* at 4 °C for two minutes and the supernatant was used as the DNA template for PCR amplification. The PCR was performed with 20 µL Mango Mix™ (Bioline), 0.25 µM of each primer and 2 µL of DNA template in a final volume of 40 µL. The thermal cycling protocol was as follows: 95 °C for 5 min; 35 cycles of 95 °C for 1 min, 50 °C for 1 min, and 72 °C for 1 min; and a final extension of 10 min at 72 °C. Amplicons were purified and Sanger sequenced on an ABI sequencing instrument by Macrogen Inc. (Seoul, South Korea) or by the Australian Genome Research Facility (AGRF) using the 1492r primer. Trimmed high quality read data from each isolate was used for presumptive identification by querying the 16S rRNA gene sequences via BLASTn. For some isolates the near-complete 16S rRNA gene sequence was determined by sequencing with additional primers (27f, 357f (5′-CCT ACG GGA GGC AGC AG-3′, (94)), 926f (5’-CCG TCA ATT CMT TTR AGT TT-3’, (95)), 519r (5’-GWA TTA CCG CGG CKG CTG-3’, (94)), 926r (5’-AAA CTR AAA MGA ATT GAC GG-3’, (95)), and 1492r). The six reads for each isolate were aligned using Geneious Prime 2019.1.2 (https://www.geneious.com) via the Geneious global alignment default settings with automatic determination of read direction. From this alignment, a consensus sequence for the 16S rRNA gene was constructed based on the frequency of a base and its quality (from chromatogram data) in each alignment column. The consensus sequence length for each of the six isolate pairs varied from 1352 to 1495 nucleotides. GenBank accession numbers for sequences are shown in Table 1.

### Qualitative free radical scavenging assay

DPPH is a stable free radical that is purple in its oxidized state but becomes white-yellow when reduced by antioxidants, and has been used to identify FRS marine bacteria (96, 97). To qualitatively assess *E. diaphana*-associated bacteria isolates for FRS ability, a sterile Whatman #1 filter paper was gently pressed against fresh (2-4 days old) colonies from a streak plate. Plates (with filter paper) were then incubated overnight at 26 °C. The following day, filter papers were removed with forceps, allowed to dry in a fume hood for 30 min, and 500 µL of a 0.2 mM DPPH (Cat# D9132, Sigma-Aldrich) solution in methanol was applied with a pipette over individual colonies. As a positive control, a few drops of 0.1% (w/v) L-ascorbic acid (Cat# A7631, Sigma-Aldrich) were placed on a separate filter. The response of each isolate to DPPH was recorded within 3 min of DPPH application; a positive response was recorded if a white-yellow halo appeared around individual colonies within 1 min, a weak positive response was assigned to strains that had a halo form between 1 and 3 min after DPPH application, and a negative response was listed for strains that failed to form a halo (Fig. 3). Approximately 700 isolates were screened using the qualitative DPPH assay.

**FIG 3:**
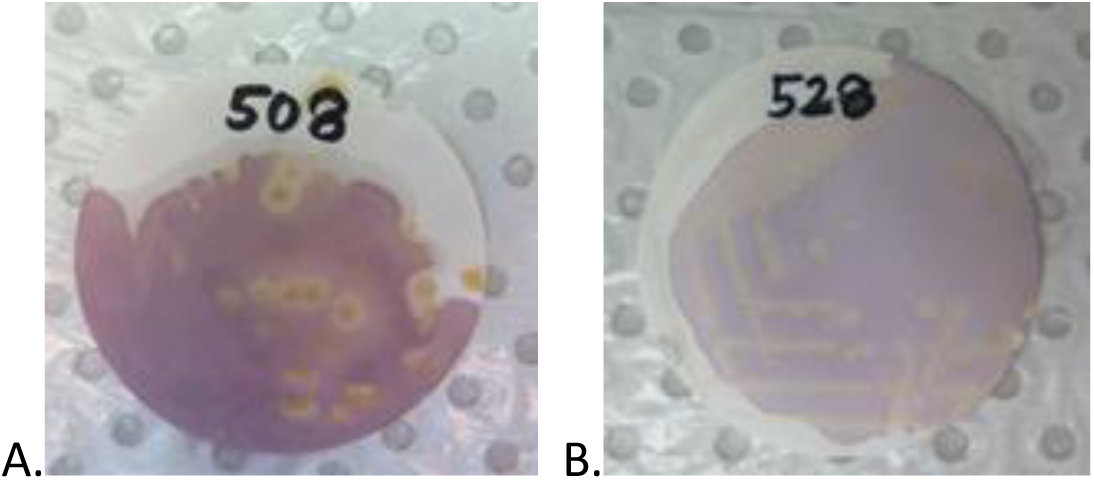
DPPH is a stable free radical that is purple in its oxidized state. When reduced by an antioxidant, a white-yellow halo will appear around individual bacteria colonies (A), this was qualitatively deemed a positive response. Isolates that did not have a halo around colonies within 3 min of DPPH application were deemed negative (B).

### Quantitative free radical scavenging assay

To quantitatively assess the FRS ability, select isolates were grown in R2A broth (Table S5). A volume of 50 mL of autoclaved medium was added to sterile 250 mL Erlenmeyer flasks and each flask was inoculated with an isolate colony grown from a R2A or MA plate culture. Cultures were grown with shaking (150 rpm; Ratek orbital incubator) at 37°C for 48 h. Sterile uninoculated R2A was used as a control. A minimum of three replicate cultures were grown per isolate. After 48 h, the optical density of each culture was measured at 600 nm (OD_600_, CLARIOstar PLUS, BMG Labtech), and the cultures

(including negative medium controls) were centrifuged at 3000 x *g* at 4°C for 30 min (Allegra X-12R) to pellet the bacterial cells. The cell-free supernatants (CFSs) were collected, frozen at – 80 °C, freeze dried (Alpha 1-4 LDplus, Martin Christ), and stored under inert gas in a dark, dry environment until analysis. Antioxidants were extracted from the CFSs by resuspending at 50 mg mL^−1^ in 100% methanol, sonicating (Branson 2510) for 5 min, then centrifuging at 3000 x *g* at 4 °C for 5 min. Quantitative DPPH assays were run by creating a 1:1 solution of 0.2 mM DPPH in methanol and CFS extract to a final volume of 1 mL, vortexing, and incubation in the dark at room temperature for 30 min. Samples were then vortexed briefly, and three 300 µL replicates of each sample were transferred into a well of a 96 well plate. Decolourization of DPPH was determined by measuring the decrease in absorbance at 517 nm (Enspire 2300 plate reader, Perkin Elmer), and the FRS activity was calculated according to the formula, % DPPH scavenging activity = (Control – Sample) / Control ×100, where, Control is the absorbance of the DPPH control (1:1 0.2 mM DPPH:methanol), and Sample is the absorbance of CFS extract in DPPH. All samples were measured against a 100% methanol blank. Positive controls consisting of 0.01 – 0.001% (w/v) L-ascorbic acid were run on each 96-well plate. FRS activity ranged from 0-90%.

### Catalase assay

The pelleted cells from above were resuspended in 2 mL fRSS and 500 µL H_2_O_2_ giving a final concentration of 16 mM. If bubbles appeared, the organism was considered catalase positive. If there were no bubbles, the organism was classified as catalase negative.

### Inhibition testing

Each paired set of high and low FRS strains were inoculated crosswise along the middle of MA plates to test for antagonism. Plates were kept at 26 °C and monitored daily for up to 7 days for antagonistic activity by documenting the presence or absence of both inoculated isolates and if there was a zone of inhibition between them.

### Phylogenetic analysis

All partial 16S rRNA gene sequences (842) were aligned with reference sequences (72) of closely related organisms using Geneious Prime 2019.1.2 (https://www.geneious.com). This alignment was used to construct a neighbor-joining phylogenetic tree using the Jukes-Cantor method. Maximum-likelihood dendrograms were generated with bootstrap values of 1000.

### Whole genome sequence analysis

Positive FRS strains along with conspecific or congeneric negative FRS strains were selected for genome sequencing; in total, six pairs of isolates were sequenced. Genomic DNA was isolated from a single colony using a JANUS Chemagic Workstation and Chemagic Viral DNA/RNA kit (PerkinElmer). Libraries were prepared with the Nextera XT DNA sample preparation kit (Illumina). Readsets were produced using the Illumina sequencing platform (Instrument: Illumina NextSeq 500, 150 base, paired-end) and the whole genome shotgun (WGS) method. Read depth coverage was approximately 100 times assuming a genome size of 4 M bases.

Illumina readsets for each isolate were assembled using Skesa (98) and the draft genome sequence annotated using Prokka (99). A genome sequence based taxonomic classification for each isolate was determined using Kraken2 (100) with the Genome Taxonomy Database (GTDB; 101) as the curated genomic data source. Classification was primarily based on the genome sequence of related isolates (within the relevant species where possible), which were obtained from GenBank. In situations where genomes of taxonomically relevant individuals were available, a species level classification was possible. Where available, closed genome sequences from GenBank were used for comparative genomics analysis. For each of the six pairs of isolates, core genome (i.e. genes shared between the isolate pair) comparisons were performed, as implemented in Nullarbor (https://github.com/tseemann/nullarbor),, with phylogenies inferred using SNP differences. Genes of interest for DMSP synthesis and degradation, vitamin B_12_ synthesis, and catalase were identified from the annotated genome sequence (GFF format) produced by Prokka; specific genes were identified by both name and Refseq accession number.

### 16S rRNA gene copy number estimation

The 16S rRNA gene copy number of the 12 draft genomes was predicted by the 16Stimator pipeline (102). Briefly, all the 12 genomic assemblies were submitted to the rapid annotations using subsystems technology (RAST) server (103), and the positions of 16S rRNA and a set of single-copied housekeeping genes (Table S6) were extracted from the RAST annotations. The clean read sets were mapped back to the corresponding genomic assemblies by Bowtie 2 (104) to determine the read depth of each position. Finally, the 16S copy number of each isolate was calculated by dividing the median depth of 16S gene by the median depth of the single-copied housekeeping genes after the read depths were calibrated by the model parameters provided by 16Stimator.

### Statistical analysis

CFU counts were analyzed in R (v3.6.2, 105) by first checking the assumptions of equal variance and homogeneity. An analysis of variance test was used to detect differences in the mean number of bacterial colonies from each anemone genotype by solid growth media (R2A or MA). A one-way analysis of variance (one-way ANOVA; 106) was used to determine if there were significant differences between FRS abilities of selected positive (high FRS), negative (low FRS), and media controls, and pairwise comparisons were performed using Tukey’s HSD (107, 108). Each probiotic pair and media control was tested to determine if data met the assumptions of normality and homoscedasticity. If either assumption was violated, the non-parametic Kruskal-Wallis rank sum test (109) was used with a Dunn test (110) for multiple comparisons (p-values adjusted with the Benjamini-Hochberg method (111)) with the R package “FSA” (112).

### Data availability

WGS raw reads are freely available in the Sequence Read Archive under BioProject PRJNA574193; the complete data set is listed in Table S1.

## ACKNOWLEDGEMENTS

This research was supported by the Australian Research Council Discovery Project grant DP160101468 (to MJHvO and LLB). MJHvO acknowledges Australian Research Council Laureate Fellowship FL180100036. We are grateful to Leon Hartman, Giulia Holland, and Shona Elliot-Kerr for their contributions in the preliminary culturing and screening of anemone-associated bacteria. Dr. Gayle Philip contributed with bacterial whole genome sequence analysis and Leon Hartman assisted with figure designs and reviewed the manuscript. Xavier Smith assisted with bacterial inhibition tests. Whole genome sequencing was organized by Dr. Glen Carter at the Peter Doherty Institute, Melbourne, Australia.

AMD, MvO and LLB conceived and designed the study. AMD performed the sampling and sample processing. AMD, DB, and HL completed bioinformatic analyses. AMD wrote the first draft. All authors edited and approved the final manuscript.

---

**Table S1:**
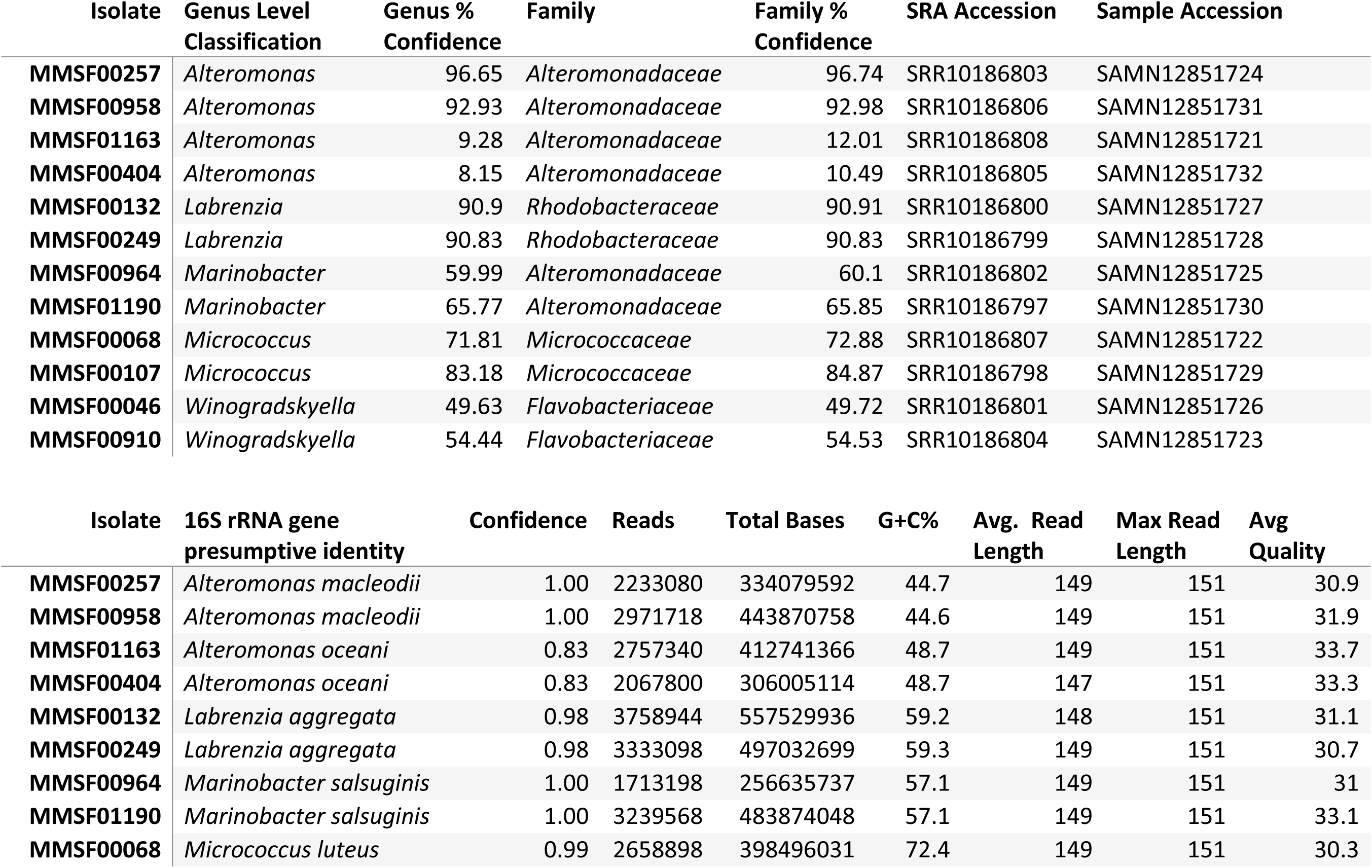
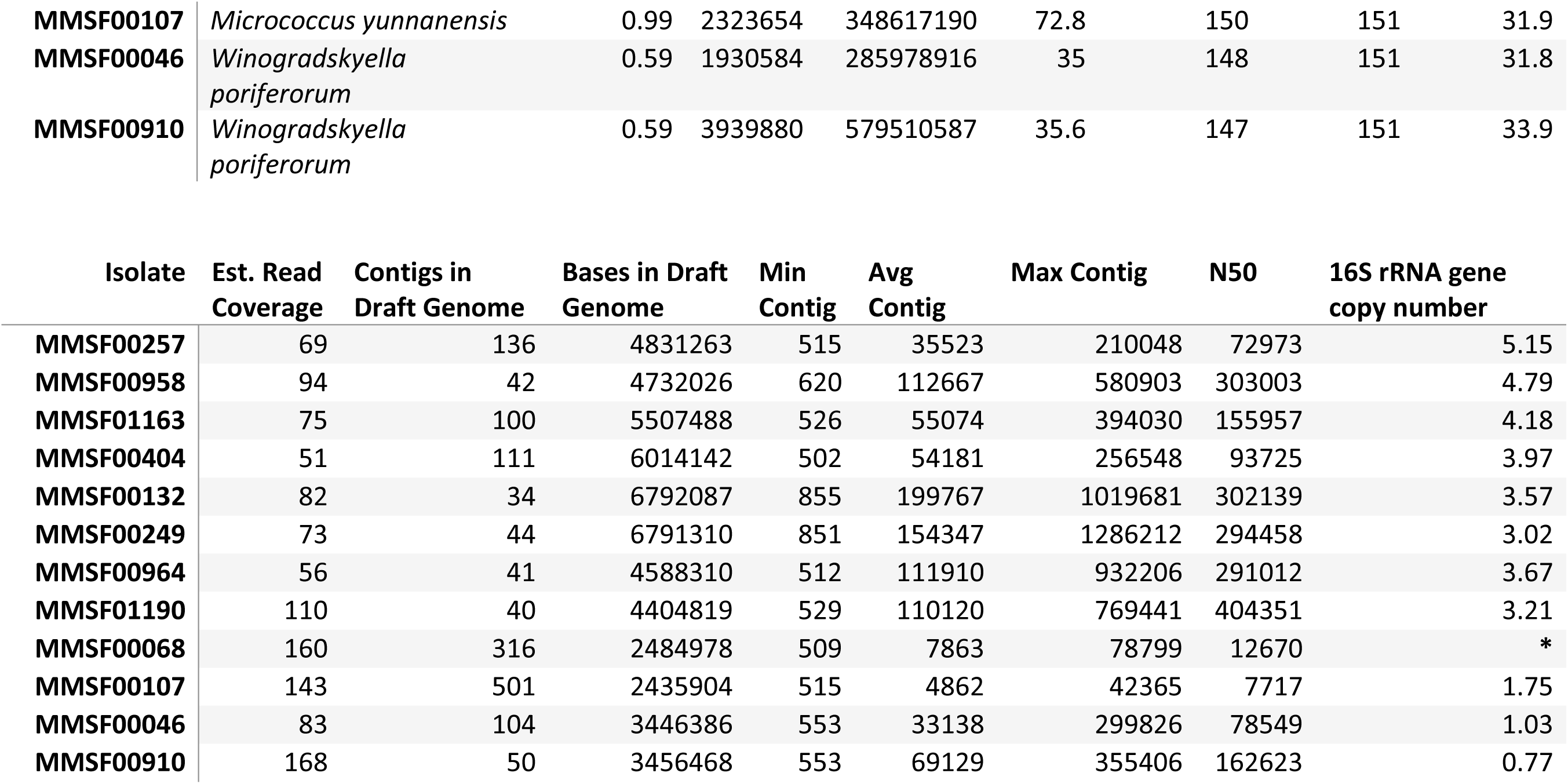
Isolate Genome Sequence Data Summary. Strains are presented as high FRS (grey) followed by low FRS (white). 16S rRNA gene presumptive identity is derived from the NCBI classification of near-complete 16S rRNA gene sequences. *We were unable to determine the 16S rRNA copy number of isolate MMSF00068 due to the presence of contaminant sequences in the WGS.

| Isolate | Genus Level Classification | Genus % Confidence | Family | Family % Confidence | SRA Accession | Sample Accession |
| --- | --- | --- | --- | --- | --- | --- |
| **MMSF00257** | *Alteromonas* | 96.65 | *Alteromonadaceae* | 96.74 | SRR10186803 | SAMN12851724 |
| **MMSF00958** | *Alteromonas* | 92.93 | *Alteromonadaceae* | 92.98 | SRR10186806 | SAMN12851731 |
| **MMSF01163** | *Alteromonas* | 9.28 | *Alteromonadaceae* | 12.01 | SRR10186808 | SAMN12851721 |
| **MMSF00404** | *Alteromonas* | 8.15 | *Alteromonadaceae* | 10.49 | SRR10186805 | SAMN12851732 |
| **MMSF00132** | *Labrenzia* | 90.9 | *Rhodobacteraceae* | 90.91 | SRR10186800 | SAMN12851727 |
| **MMSF00249** | *Labrenzia* | 90.83 | *Rhodobacteraceae* | 90.83 | SRR10186799 | SAMN12851728 |
| **MMSF00964** | *Marinobacter* | 59.99 | *Alteromonadaceae* | 60.1 | SRR10186802 | SAMN12851725 |
| **MMSF01190** | *Marinobacter* | 65.77 | *Alteromonadaceae* | 65.85 | SRR10186797 | SAMN12851730 |
| **MMSF00068** | *Micrococcus* | 71.81 | *Micrococcaceae* | 72.88 | SRR10186807 | SAMN12851722 |
| **MMSF00107** | *Micrococcus* | 83.18 | *Micrococcaceae* | 84.87 | SRR10186798 | SAMN12851729 |
| **MMSF00046** | *Winogradskyella* | 49.63 | *Flavobacteriaceae* | 49.72 | SRR10186801 | SAMN12851726 |
| **MMSF00910** | *Winogradskyella* | 54.44 | *Flavobacteriaceae* | 54.53 | SRR10186804 | SAMN12851723 |

| Isolate | 16S rRNA gene presumptive identity | Confidence | Reads | Total Bases | G+C% | Avg. Read Length | Max Read Length | Avg Quality |
| --- | --- | --- | --- | --- | --- | --- | --- | --- |
| **MMSF00257** | *Alteromonas macleodii* | 1.00 | 2233080 | 334079592 | 44.7 | 149 | 151 | 30.9 |
| **MMSF00958** | *Alteromonas macleodii* | 1.00 | 2971718 | 443870758 | 44.6 | 149 | 151 | 31.9 |
| **MMSF01163** | *Alteromonas oceani* | 0.83 | 2757340 | 412741366 | 48.7 | 149 | 151 | 33.7 |
| **MMSF00404** | *Alteromonas oceani* | 0.83 | 2067800 | 306005114 | 48.7 | 147 | 151 | 33.3 |
| **MMSF00132** | *Labrenzia aggregata* | 0.98 | 3758944 | 557529936 | 59.2 | 148 | 151 | 31.1 |
| **MMSF00249** | *Labrenzia aggregata* | 0.98 | 3333098 | 497032699 | 59.3 | 149 | 151 | 30.7 |
| **MMSF00964** | *Marinobacter salsuginis* | 1.00 | 1713198 | 256635737 | 57.1 | 149 | 151 | 31 |
| **MMSF01190** | *Marinobacter salsuginis* | 1.00 | 3239568 | 483874048 | 57.1 | 149 | 151 | 33.1 |
| **MMSF00068** | *Micrococcus luteus* | 0.99 | 2658898 | 398496031 | 72.4 | 149 | 151 | 30.3 |

|  |  |  |  |  |  |  |  |  |
| --- | --- | --- | --- | --- | --- | --- | --- | --- |
| **MMSF00107** | *Micrococcus yunnanensis* | 0.99 | 2323654 | 348617190 | 72.8 | 150 | 151 | 31.9 |
| **MMSF00046** | *Winogradskyella poriferorum* | 0.59 | 1930584 | 285978916 | 35 | 148 | 151 | 31.8 |
| **MMSF00910** | *Winogradskyella poriferorum* | 0.59 | 3939880 | 579510587 | 35.6 | 147 | 151 | 33.9 |

| Isolate | Est. Read Coverage | Contigs in Draft Genome | Bases in Draft Genome | Min Contig | Avg Contig | Max Contig | N50 | 16S rRNA gene copy number |
| --- | --- | --- | --- | --- | --- | --- | --- | --- |
| **MMSF00257** | 69 | 136 | 4831263 | 515 | 35523 | 210048 | 72973 | 5.15 |
| **MMSF00958** | 94 | 42 | 4732026 | 620 | 112667 | 580903 | 303003 | 4.79 |
| **MMSF01163** | 75 | 100 | 5507488 | 526 | 55074 | 394030 | 155957 | 4.18 |
| **MMSF00404** | 51 | 111 | 6014142 | 502 | 54181 | 256548 | 93725 | 3.97 |
| **MMSF00132** | 82 | 34 | 6792087 | 855 | 199767 | 1019681 | 302139 | 3.57 |
| **MMSF00249** | 73 | 44 | 6791310 | 851 | 154347 | 1286212 | 294458 | 3.02 |
| **MMSF00964** | 56 | 41 | 4588310 | 512 | 111910 | 932206 | 291012 | 3.67 |
| **MMSF01190** | 110 | 40 | 4404819 | 529 | 110120 | 769441 | 404351 | 3.21 |
| **MMSF00068** | 160 | 316 | 2484978 | 509 | 7863 | 78799 | 12670 | * |
| **MMSF00107** | 143 | 501 | 2435904 | 515 | 4862 | 42365 | 7717 | 1.75 |
| **MMSF00046** | 83 | 104 | 3446386 | 553 | 33138 | 299826 | 78549 | 1.03 |
| **MMSF00910** | 168 | 50 | 3456468 | 553 | 69129 | 355406 | 162623 | 0.77 |

**Table S2:** Pairwise comparison of the genome sequences between the pairs of isolates

| NCBI Classification | Type strain | Core Genome | SNP difference |
| --- | --- | --- | --- |
| *Alteromonas macleodii* | ATCC 27126 | 85% | ~60000 |
| *Alteromonas oceani* | S35 | 60% | ~190000 |
| *Labrenzia aggregata* | IAM12614 | 55% | 5 |
| *Marinobacter salsuginis* | SD14B | 15% | ~120000 |
| *Micrococcus* spp. | NCTC2665 | 70% | ~35000 |
| *Winogradskyella poriferorum* | NA | NA | 10 |

**Table S3:**
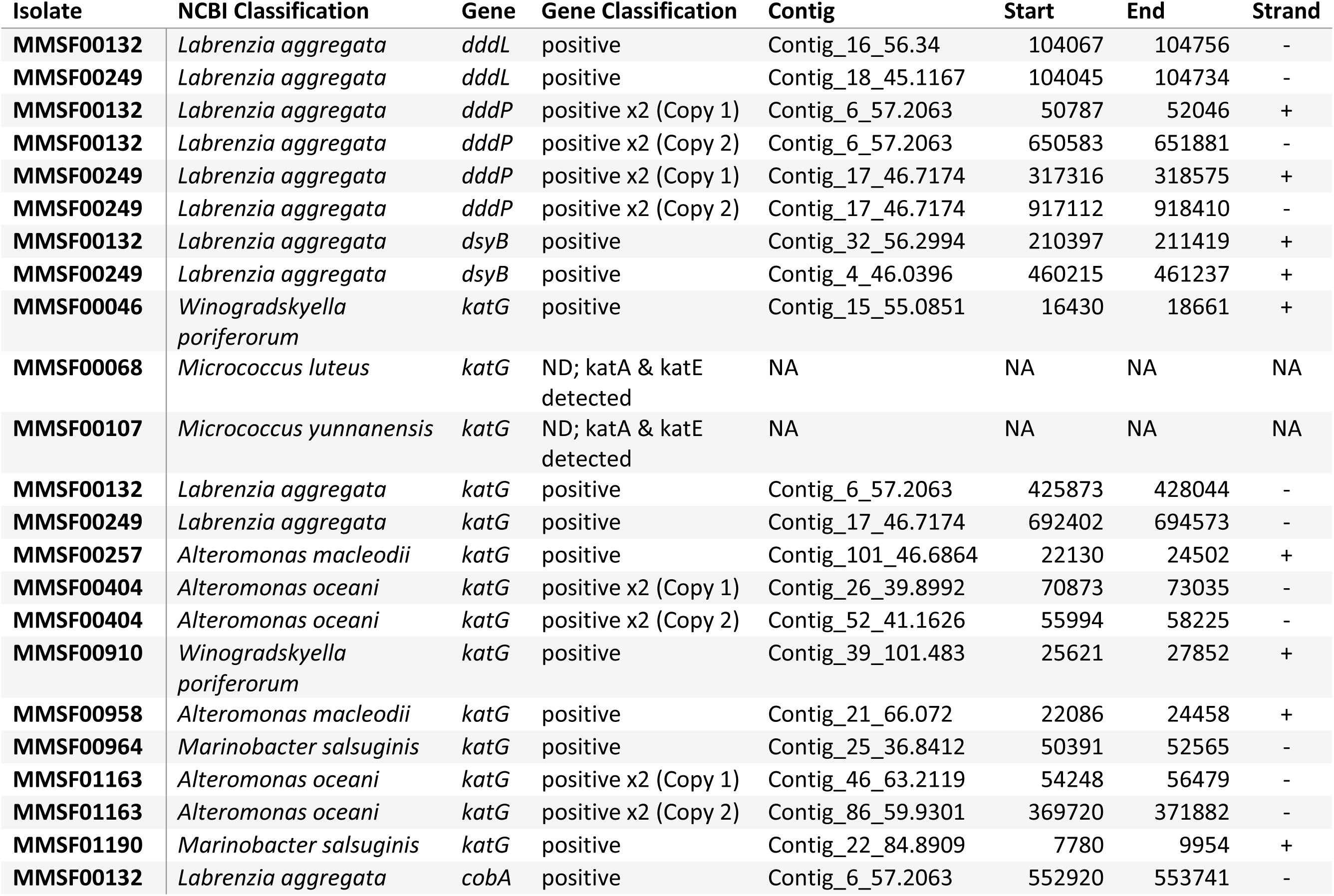
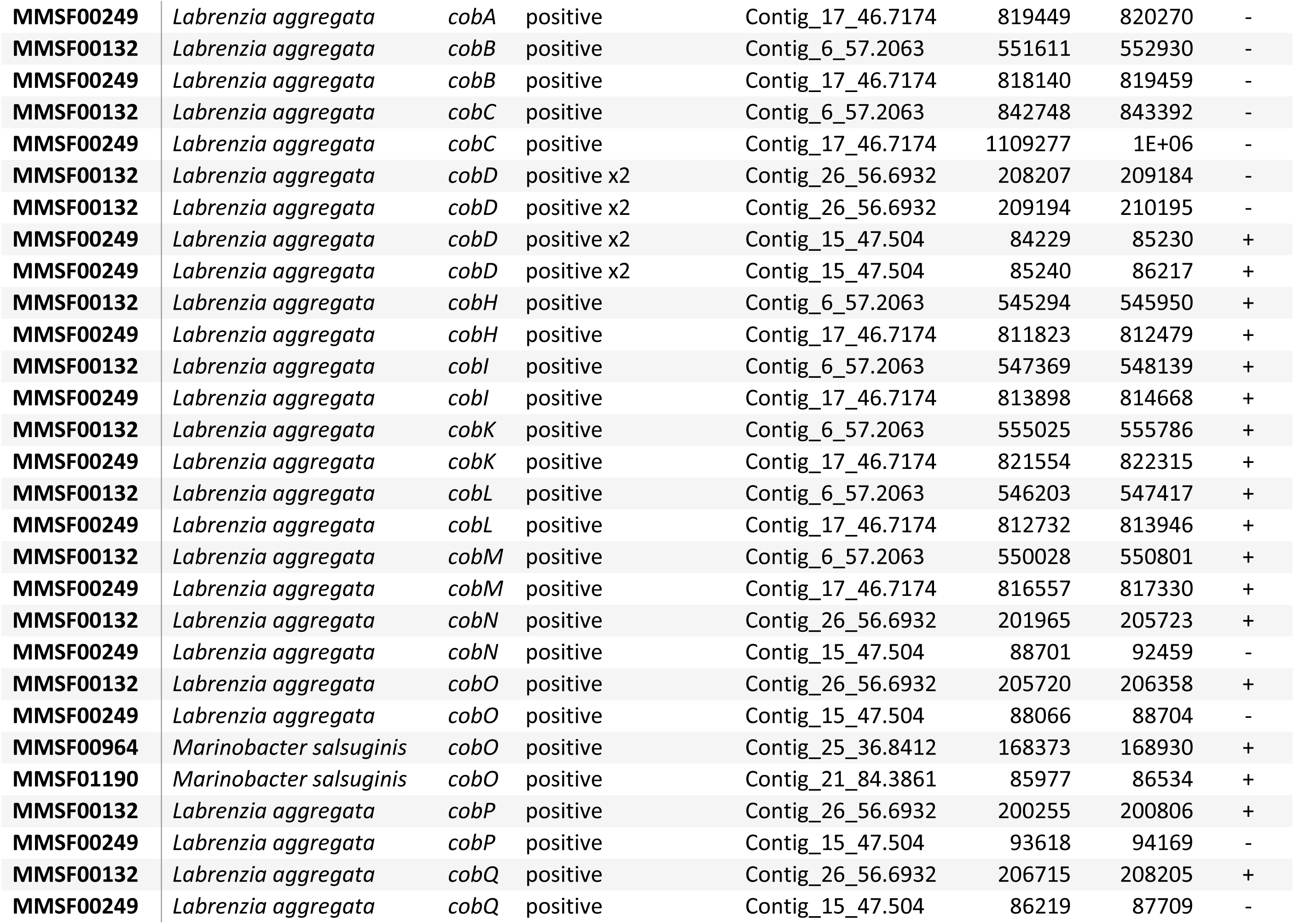
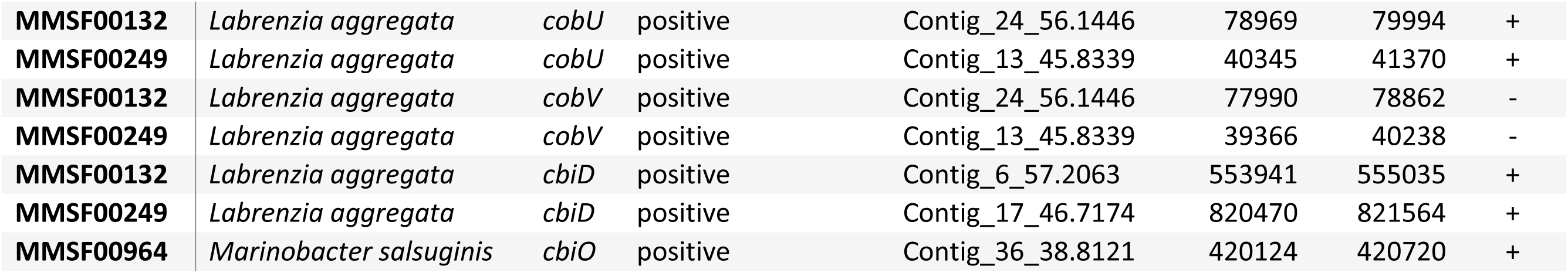
Search outcomes for genes of interest.

| Isolate | NCBI Classification | Gene | Gene Classification | Contig | Start | End | Strand |
| --- | --- | --- | --- | --- | --- | --- | --- |
| MMSF00132 | *Labrenzia aggregata* | *dddL* | positive | Contig_16_56.34 | 104067 | 104756 | - |
| MMSF00249 | *Labrenzia aggregata* | *dddL* | positive | Contig_18_45.1167 | 104045 | 104734 | - |
| MMSF00132 | *Labrenzia aggregata* | *dddP* | positive x2 (Copy 1) | Contig_6_57.2063 | 50787 | 52046 | + |
| MMSF00132 | *Labrenzia aggregata* | *dddP* | positive x2 (Copy 2) | Contig_6_57.2063 | 650583 | 651881 | - |
| MMSF00249 | *Labrenzia aggregata* | *dddP* | positive x2 (Copy 1) | Contig_17_46.7174 | 317316 | 318575 | + |
| MMSF00249 | *Labrenzia aggregata* | *dddP* | positive x2 (Copy 2) | Contig_17_46.7174 | 917112 | 918410 | - |
| MMSF00132 | *Labrenzia aggregata* | *dsyB* | positive | Contig_32_56.2994 | 210397 | 211419 | + |
| MMSF00249 | *Labrenzia aggregata* | *dsyB* | positive | Contig_4_46.0396 | 460215 | 461237 | + |
| MMSF00046 | *Winogradskyella*<br>*poriferorum* | *katG* | positive | Contig_15_55.0851 | 16430 | 18661 | + |
| MMSF00068 | *Micrococcus luteus* | *katG* | ND; katA & katE<br>detected | NA | NA | NA | NA |
| MMSF00107 | *Micrococcus yunnanensis* | *katG* | ND; katA & katE<br>detected | NA | NA | NA | NA |
| MMSF00132 | *Labrenzia aggregata* | *katG* | positive | Contig_6_57.2063 | 425873 | 428044 | - |
| MMSF00249 | *Labrenzia aggregata* | *katG* | positive | Contig_17_46.7174 | 692402 | 694573 | - |
| MMSF00257 | *Alteromonas macleodii* | *katG* | positive | Contig_101_46.6864 | 22130 | 24502 | + |
| MMSF00404 | *Alteromonas oceani* | *katG* | positive x2 (Copy 1) | Contig_26_39.8992 | 70873 | 73035 | - |
| MMSF00404 | *Alteromonas oceani* | *katG* | positive x2 (Copy 2) | Contig_52_41.1626 | 55994 | 58225 | - |
| MMSF00910 | *Winogradskyella*<br>*poriferorum* | *katG* | positive | Contig_39_101.483 | 25621 | 27852 | + |
| MMSF00958 | *Alteromonas macleodii* | *katG* | positive | Contig_21_66.072 | 22086 | 24458 | + |
| MMSF00964 | *Marinobacter salsuginis* | *katG* | positive | Contig_25_36.8412 | 50391 | 52565 | - |
| MMSF01163 | *Alteromonas oceani* | *katG* | positive x2 (Copy 1) | Contig_46_63.2119 | 54248 | 56479 | - |
| MMSF01163 | *Alteromonas oceani* | *katG* | positive x2 (Copy 2) | Contig_86_59.9301 | 369720 | 371882 | - |
| MMSF01190 | *Marinobacter salsuginis* | *katG* | positive | Contig_22_84.8909 | 7780 | 9954 | + |
| MMSF00132 | *Labrenzia aggregata* | *cobA* | positive | Contig_6_57.2063 | 552920 | 553741 | - |
| **MMSF00249** | *Labrenzia aggregata* | *cobA* | positive | Contig_17_46.7174 | 819449 | 820270 | - |
| **MMSF00132** | *Labrenzia aggregata* | *cobB* | positive | Contig_6_57.2063 | 551611 | 552930 | - |
| **MMSF00249** | *Labrenzia aggregata* | *cobB* | positive | Contig_17_46.7174 | 818140 | 819459 | - |
| **MMSF00132** | *Labrenzia aggregata* | *cobC* | positive | Contig_6_57.2063 | 842748 | 843392 | - |
| **MMSF00249** | *Labrenzia aggregata* | *cobC* | positive | Contig_17_46.7174 | 1109277 | 1E+06 | - |
| **MMSF00132** | *Labrenzia aggregata* | *cobD* | positive x2 | Contig_26_56.6932 | 208207 | 209184 | - |
| **MMSF00132** | *Labrenzia aggregata* | *cobD* | positive x2 | Contig_26_56.6932 | 209194 | 210195 | - |
| **MMSF00249** | *Labrenzia aggregata* | *cobD* | positive x2 | Contig_15_47.504 | 84229 | 85230 | + |
| **MMSF00249** | *Labrenzia aggregata* | *cobD* | positive x2 | Contig_15_47.504 | 85240 | 86217 | + |
| **MMSF00132** | *Labrenzia aggregata* | *cobH* | positive | Contig_6_57.2063 | 545294 | 545950 | + |
| **MMSF00249** | *Labrenzia aggregata* | *cobH* | positive | Contig_17_46.7174 | 811823 | 812479 | + |
| **MMSF00132** | *Labrenzia aggregata* | *cobI* | positive | Contig_6_57.2063 | 547369 | 548139 | + |
| **MMSF00249** | *Labrenzia aggregata* | *cobI* | positive | Contig_17_46.7174 | 813898 | 814668 | + |
| **MMSF00132** | *Labrenzia aggregata* | *cobK* | positive | Contig_6_57.2063 | 555025 | 555786 | + |
| **MMSF00249** | *Labrenzia aggregata* | *cobK* | positive | Contig_17_46.7174 | 821554 | 822315 | + |
| **MMSF00132** | *Labrenzia aggregata* | *cobL* | positive | Contig_6_57.2063 | 546203 | 547417 | + |
| **MMSF00249** | *Labrenzia aggregata* | *cobL* | positive | Contig_17_46.7174 | 812732 | 813946 | + |
| **MMSF00132** | *Labrenzia aggregata* | *cobM* | positive | Contig_6_57.2063 | 550028 | 550801 | + |
| **MMSF00249** | *Labrenzia aggregata* | *cobM* | positive | Contig_17_46.7174 | 816557 | 817330 | + |
| **MMSF00132** | *Labrenzia aggregata* | *cobN* | positive | Contig_26_56.6932 | 201965 | 205723 | + |
| **MMSF00249** | *Labrenzia aggregata* | *cobN* | positive | Contig_15_47.504 | 88701 | 92459 | - |
| **MMSF00132** | *Labrenzia aggregata* | *cobO* | positive | Contig_26_56.6932 | 205720 | 206358 | + |
| **MMSF00249** | *Labrenzia aggregata* | *cobO* | positive | Contig_15_47.504 | 88066 | 88704 | - |
| **MMSF00964** | *Marinobacter salsuginis* | *cobO* | positive | Contig_25_36.8412 | 168373 | 168930 | + |
| **MMSF01190** | *Marinobacter salsuginis* | *cobO* | positive | Contig_21_84.3861 | 85977 | 86534 | + |
| **MMSF00132** | *Labrenzia aggregata* | *cobP* | positive | Contig_26_56.6932 | 200255 | 200806 | + |
| **MMSF00249** | *Labrenzia aggregata* | *cobP* | positive | Contig_15_47.504 | 93618 | 94169 | - |
| **MMSF00132** | *Labrenzia aggregata* | *cobQ* | positive | Contig_26_56.6932 | 206715 | 208205 | + |
| **MMSF00249** | *Labrenzia aggregata* | *cobQ* | positive | Contig_15_47.504 | 86219 | 87709 | - |
| **MMSF00132** | *Labrenzia aggregata* | *cobU* | positive | Contig_24_56.1446 | 78969 | 79994 | + |
| **MMSF00249** | *Labrenzia aggregata* | *cobU* | positive | Contig_13_45.8339 | 40345 | 41370 | + |
| **MMSF00132** | *Labrenzia aggregata* | *cobV* | positive | Contig_24_56.1446 | 77990 | 78862 | - |
| **MMSF00249** | *Labrenzia aggregata* | *cobV* | positive | Contig_13_45.8339 | 39366 | 40238 | - |
| **MMSF00132** | *Labrenzia aggregata* | *cbiD* | positive | Contig_6_57.2063 | 553941 | 555035 | + |
| **MMSF00249** | *Labrenzia aggregata* | *cbiD* | positive | Contig_17_46.7174 | 820470 | 821564 | + |
| **MMSF00964** | *Marinobacter salsuginis* | *cbiO* | positive | Contig_36_38.8121 | 420124 | 420720 | + |

**Table S4:**
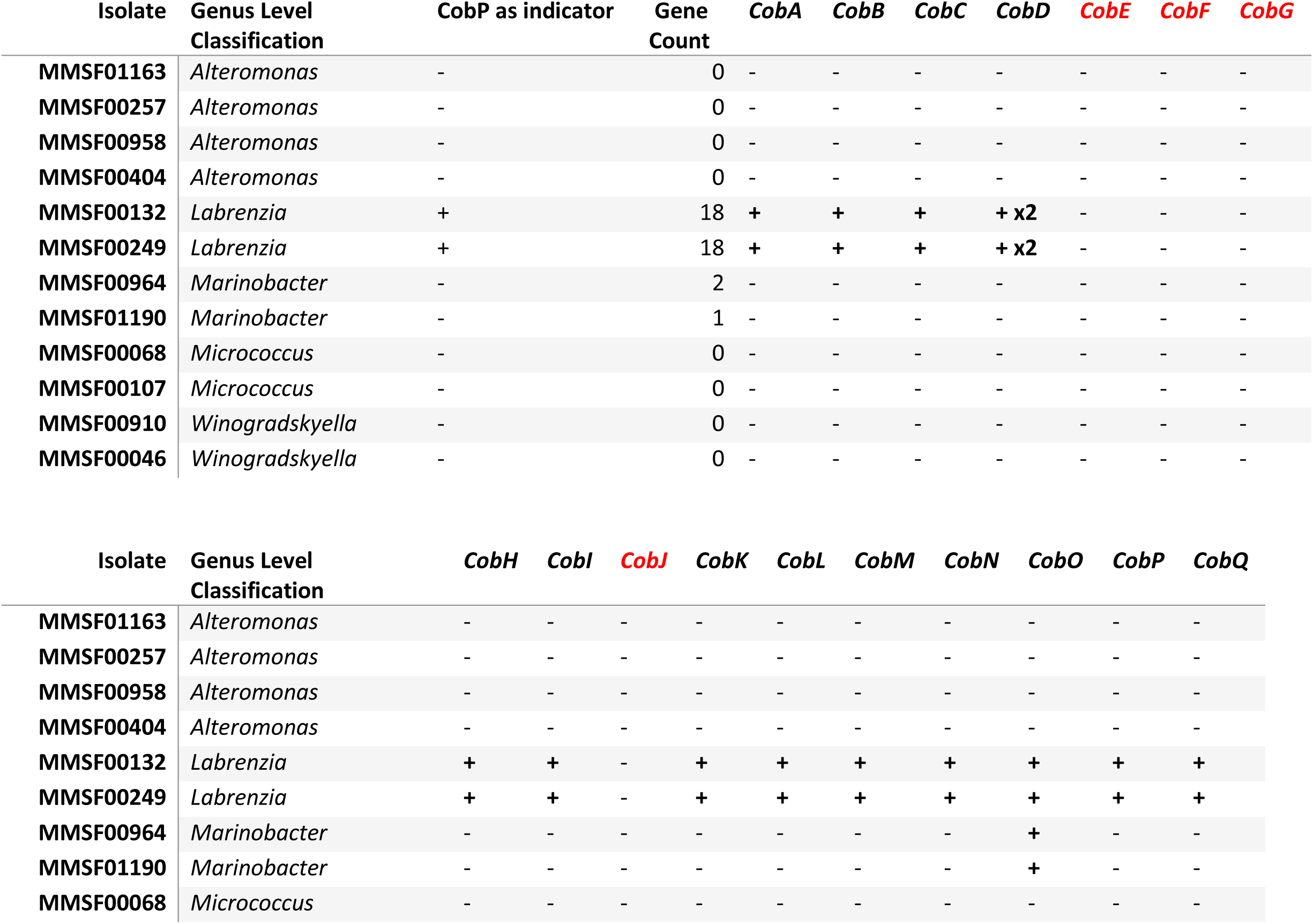
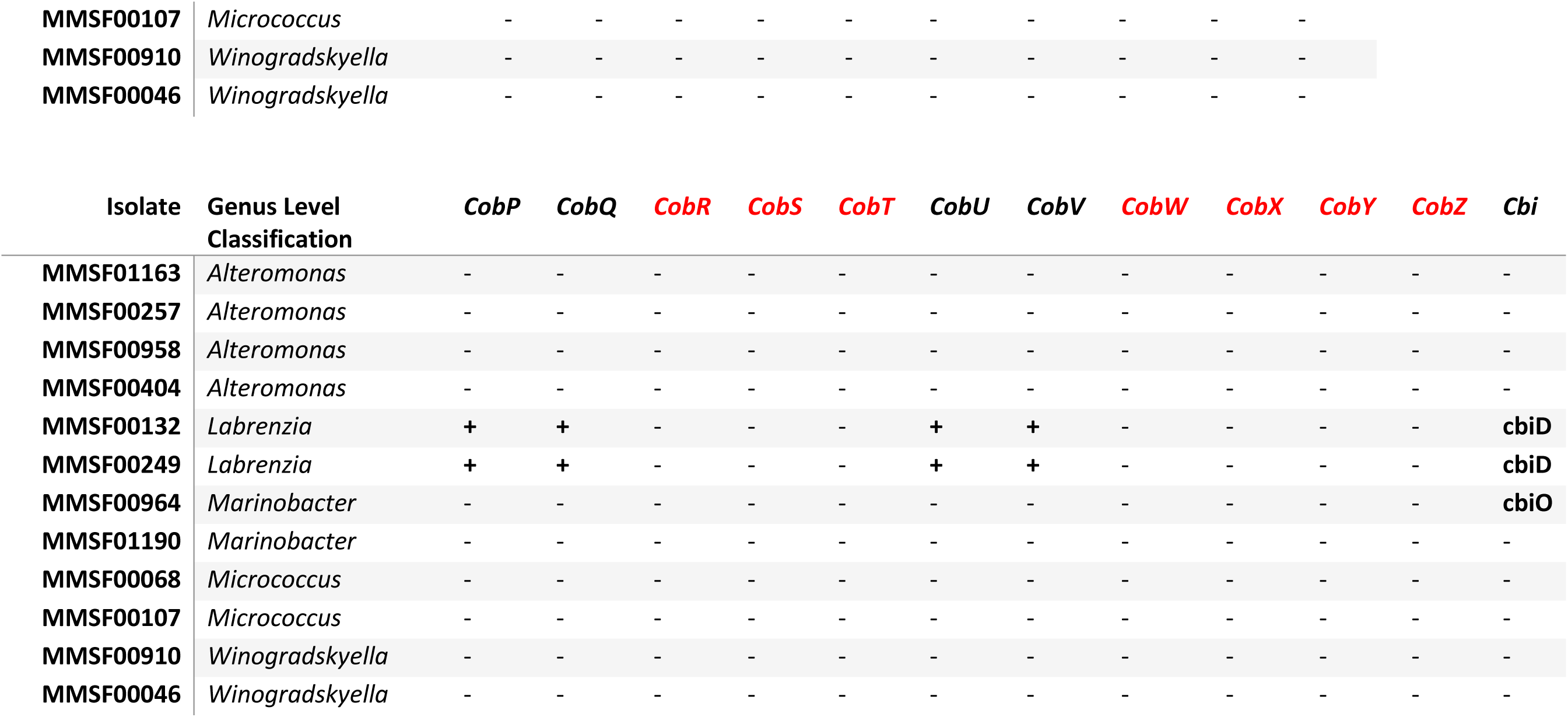
Summary of vitamin B_12_ biosynthesis pathway genes. A “+” indicates the presence of the gene in the respective isolate, whereas a “-” represents the absence of that gene. Genes in red were not found in any isolate.

**Table S5:** Composition of R2A broth adjusted to suit marine bacteria. Final pH = 7.2 +/−0.2 at 26 °C. R2A broth was made by suspending 43.12 g of combined reagents in 1 L of MilliQ water, dissolving the medium completely, and sterilization by autoclaving at 121°C for 15 min.

| Component | grams L<sup>-1</sup> | Supplier |
| --- | --- | --- |
| Casein acid hydrolysate | 0.500 | Cat#C0501, Sigma-Aldrich |
| Yeast extract | 0.500 | Cat#LP0021, Oxoid |
| Proteose peptone | 0.500 | Cat#211684, ThermoFisher |
| Dextrose | 0.500 | Cat#G360, PhytoTech Laboratories |
| Starch, soluble | 0.500 | Cat#AJA526, Univar |
| Dipotassium phosphate | 0.300 | Cat#P3786, Sigma-Aldrich |
| Magnesium sulfate | 0.024 | Cat#M2643, Sigma-Aldrich |
| Sodium pyruvate | 0.300 | Cat#P2256, Sigma-Aldrich |
| Red Sea Salt™ | 40.00 | Cat#R11065, Red Sea |

**Table S6:**
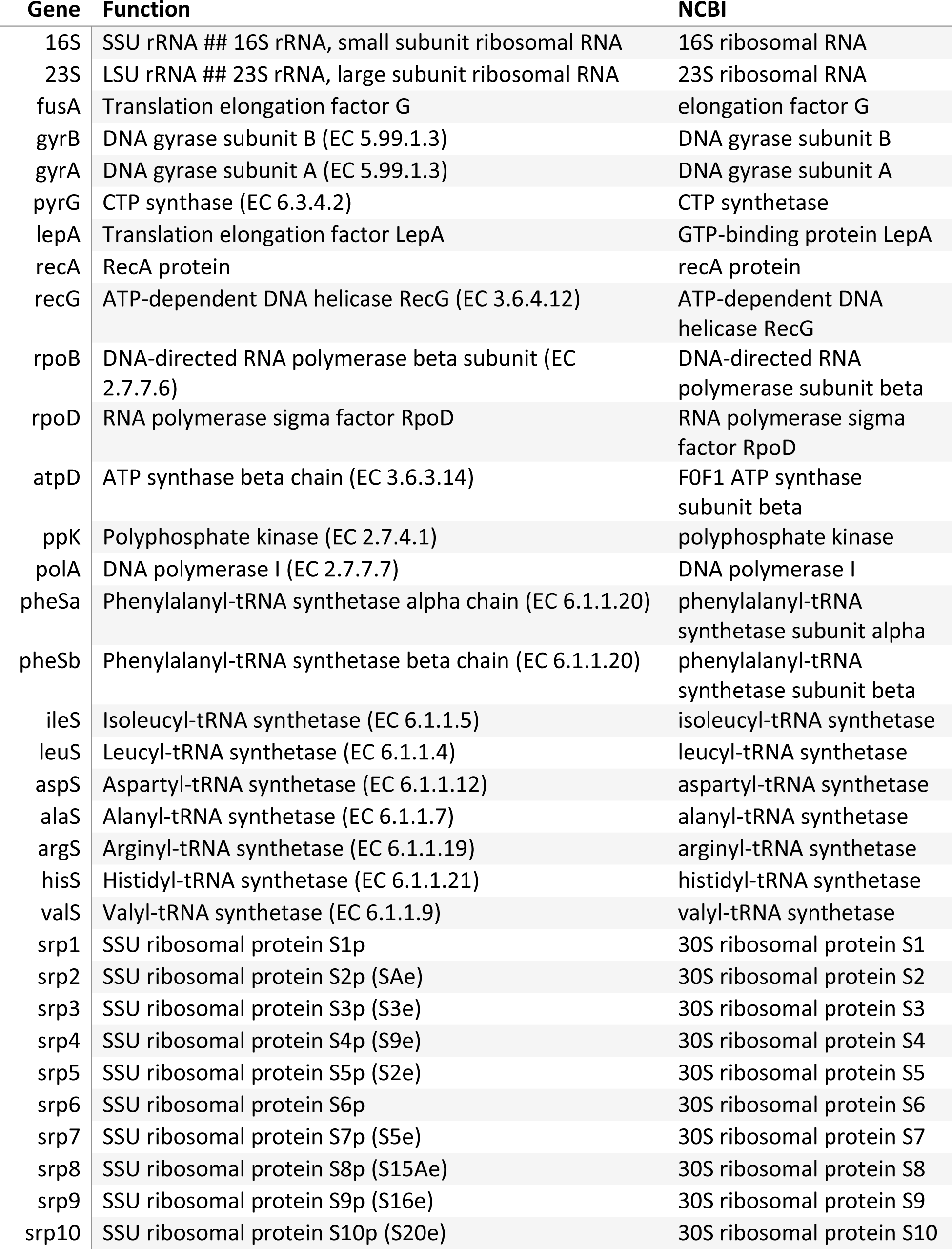
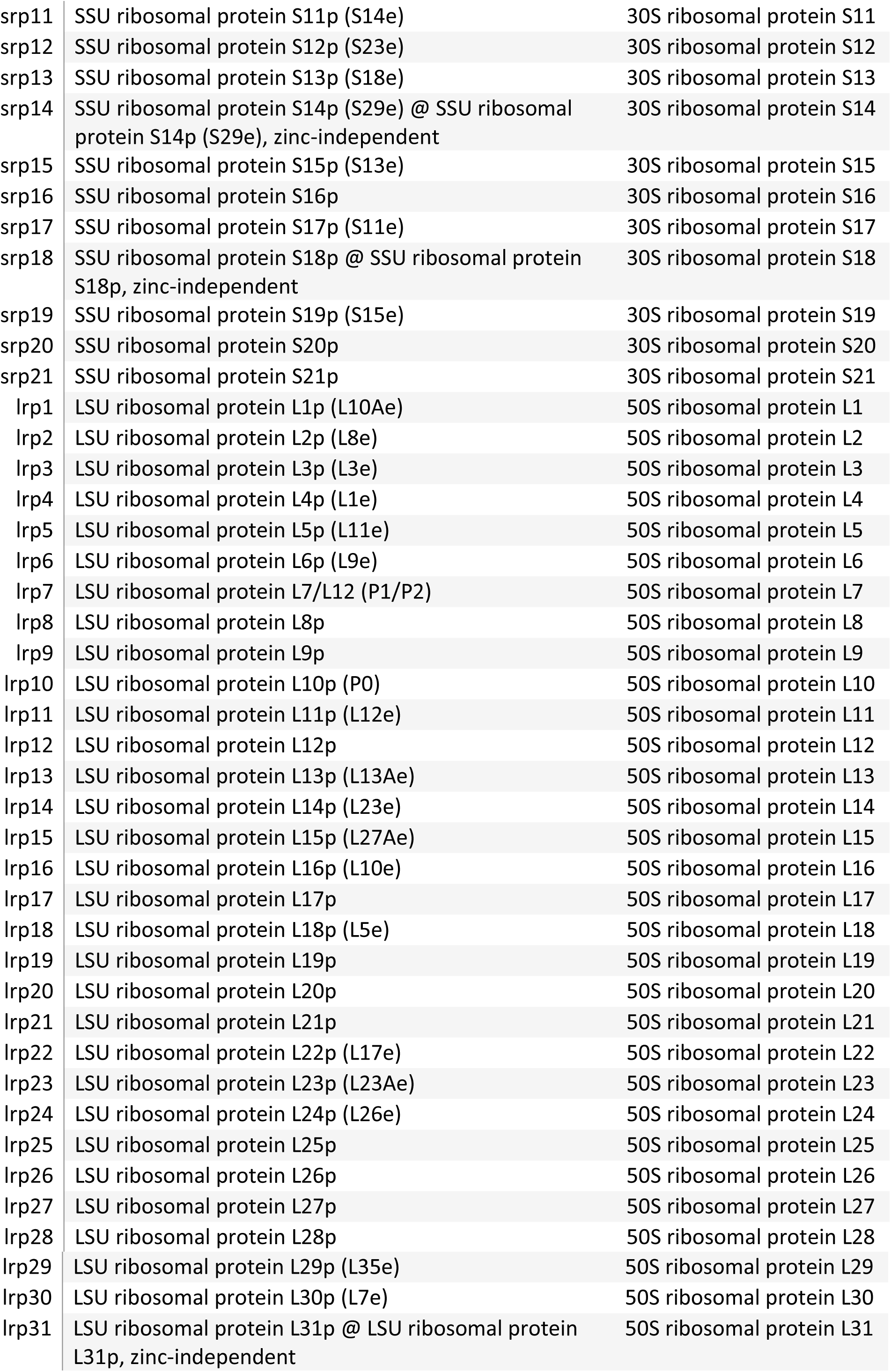
Single-copy housekeeping genes extracted from the RAST annotations

| Gene | Function | NCBI |
| --- | --- | --- |
| 16S | SSU rRNA ## 16S rRNA, small subunit ribosomal RNA | 16S ribosomal RNA |
| 23S | LSU rRNA ## 23S rRNA, large subunit ribosomal RNA | 23S ribosomal RNA |
| fusA | Translation elongation factor G | elongation factor G |
| gyrB | DNA gyrase subunit B (EC 5.99.1.3) | DNA gyrase subunit B |
| gyrA | DNA gyrase subunit A (EC 5.99.1.3) | DNA gyrase subunit A |
| pyrG | CTP synthase (EC 6.3.4.2) | CTP synthetase |
| lepA | Translation elongation factor LepA | GTP-binding protein LepA |
| recA | RecA protein | recA protein |
| recG | ATP-dependent DNA helicase RecG (EC 3.6.4.12) | ATP-dependent DNA helicase RecG |
| rpoB | DNA-directed RNA polymerase beta subunit (EC 2.7.7.6) | DNA-directed RNA polymerase subunit beta |
| rpoD | RNA polymerase sigma factor RpoD | RNA polymerase sigma factor RpoD |
| atpD | ATP synthase beta chain (EC 3.6.3.14) | FOF1 ATP synthase subunit beta |
| ppk | Polyphosphate kinase (EC 2.7.4.1) | polyphosphate kinase |
| polA | DNA polymerase I (EC 2.7.7.7) | DNA polymerase I |
| pheSa | Phenylalanyl-tRNA synthetase alpha chain (EC 6.1.1.20) | phenylalanyl-tRNA synthetase subunit alpha |
| pheSb | Phenylalanyl-tRNA synthetase beta chain (EC 6.1.1.20) | phenylalanyl-tRNA synthetase subunit beta |
| ileS | Isoleucyl-tRNA synthetase (EC 6.1.1.5) | isoleucyl-tRNA synthetase |
| leuS | Leucyl-tRNA synthetase (EC 6.1.1.4) | leucyl-tRNA synthetase |
| aspS | Aspartyl-tRNA synthetase (EC 6.1.1.12) | aspartyl-tRNA synthetase |
| alaS | Alanyl-tRNA synthetase (EC 6.1.1.7) | alanyl-tRNA synthetase |
| argS | Arginyl-tRNA synthetase (EC 6.1.1.19) | arginyl-tRNA synthetase |
| hisS | Histidyl-tRNA synthetase (EC 6.1.1.21) | histidyl-tRNA synthetase |
| valS | Valyl-tRNA synthetase (EC 6.1.1.9) | valyl-tRNA synthetase |
| srp1 | SSU ribosomal protein S1p | 30S ribosomal protein S1 |
| srp2 | SSU ribosomal protein S2p (SAe) | 30S ribosomal protein S2 |
| srp3 | SSU ribosomal protein S3p (S3e) | 30S ribosomal protein S3 |
| srp4 | SSU ribosomal protein S4p (S9e) | 30S ribosomal protein S4 |
| srp5 | SSU ribosomal protein S5p (S2e) | 30S ribosomal protein S5 |
| srp6 | SSU ribosomal protein S6p | 30S ribosomal protein S6 |
| srp7 | SSU ribosomal protein S7p (S5e) | 30S ribosomal protein S7 |
| srp8 | SSU ribosomal protein S8p (S15Ae) | 30S ribosomal protein S8 |
| srp9 | SSU ribosomal protein S9p (S16e) | 30S ribosomal protein S9 |
| srp10 | SSU ribosomal protein S10p (S20e) | 30S ribosomal protein S10 |
| srp11 | SSU ribosomal protein S11p (S14e) | 30S ribosomal protein S11 |
| srp12 | SSU ribosomal protein S12p (S23e) | 30S ribosomal protein S12 |
| srp13 | SSU ribosomal protein S13p (S18e) | 30S ribosomal protein S13 |
| srp14 | SSU ribosomal protein S14p (S29e) @ SSU ribosomal protein S14p (S29e), zinc-independent | 30S ribosomal protein S14 |
| srp15 | SSU ribosomal protein S15p (S13e) | 30S ribosomal protein S15 |
| srp16 | SSU ribosomal protein S16p | 30S ribosomal protein S16 |
| srp17 | SSU ribosomal protein S17p (S11e) | 30S ribosomal protein S17 |
| srp18 | SSU ribosomal protein S18p @ SSU ribosomal protein S18p, zinc-independent | 30S ribosomal protein S18 |
| srp19 | SSU ribosomal protein S19p (S15e) | 30S ribosomal protein S19 |
| srp20 | SSU ribosomal protein S20p | 30S ribosomal protein S20 |
| srp21 | SSU ribosomal protein S21p | 30S ribosomal protein S21 |
| lrp1 | LSU ribosomal protein L1p (L10Ae) | 50S ribosomal protein L1 |
| lrp2 | LSU ribosomal protein L2p (L8e) | 50S ribosomal protein L2 |
| lrp3 | LSU ribosomal protein L3p (L3e) | 50S ribosomal protein L3 |
| lrp4 | LSU ribosomal protein L4p (L1e) | 50S ribosomal protein L4 |
| lrp5 | LSU ribosomal protein L5p (L11e) | 50S ribosomal protein L5 |
| lrp6 | LSU ribosomal protein L6p (L9e) | 50S ribosomal protein L6 |
| lrp7 | LSU ribosomal protein L7/L12 (P1/P2) | 50S ribosomal protein L7 |
| lrp8 | LSU ribosomal protein L8p | 50S ribosomal protein L8 |
| lrp9 | LSU ribosomal protein L9p | 50S ribosomal protein L9 |
| lrp10 | LSU ribosomal protein L10p (P0) | 50S ribosomal protein L10 |
| lrp11 | LSU ribosomal protein L11p (L12e) | 50S ribosomal protein L11 |
| lrp12 | LSU ribosomal protein L12p | 50S ribosomal protein L12 |
| lrp13 | LSU ribosomal protein L13p (L13Ae) | 50S ribosomal protein L13 |
| lrp14 | LSU ribosomal protein L14p (L23e) | 50S ribosomal protein L14 |
| lrp15 | LSU ribosomal protein L15p (L27Ae) | 50S ribosomal protein L15 |
| lrp16 | LSU ribosomal protein L16p (L10e) | 50S ribosomal protein L16 |
| lrp17 | LSU ribosomal protein L17p | 50S ribosomal protein L17 |
| lrp18 | LSU ribosomal protein L18p (L5e) | 50S ribosomal protein L18 |
| lrp19 | LSU ribosomal protein L19p | 50S ribosomal protein L19 |
| lrp20 | LSU ribosomal protein L20p | 50S ribosomal protein L20 |
| lrp21 | LSU ribosomal protein L21p | 50S ribosomal protein L21 |
| lrp22 | LSU ribosomal protein L22p (L17e) | 50S ribosomal protein L22 |
| lrp23 | LSU ribosomal protein L23p (L23Ae) | 50S ribosomal protein L23 |
| lrp24 | LSU ribosomal protein L24p (L26e) | 50S ribosomal protein L24 |
| lrp25 | LSU ribosomal protein L25p | 50S ribosomal protein L25 |
| lrp26 | LSU ribosomal protein L26p | 50S ribosomal protein L26 |
| lrp27 | LSU ribosomal protein L27p | 50S ribosomal protein L27 |
| lrp28 | LSU ribosomal protein L28p | 50S ribosomal protein L28 |
| lrp29 | LSU ribosomal protein L29p (L35e) | 50S ribosomal protein L29 |
| lrp30 | LSU ribosomal protein L30p (L7e) | 50S ribosomal protein L30 |
| lrp31 | LSU ribosomal protein L31p @ LSU ribosomal protein L31p, zinc-independent | 50S ribosomal protein L31 |

